# Genetic evaluation modeling of the Tunisian dairy cattle: assessment of the test-day model (TDM) and lactation model (L305) in accordance to the flock size

**DOI:** 10.1101/589523

**Authors:** Yosra Ressaissi, Mohamed Ben Hmouda

**Affiliations:** Departement des Sciences Animales, Institut National Agronomique de Tunisie, 43 Avenue Charle Nicolle 1082-Tunis-Mahrajène-Tunisie; Institut National de recherches agronomiques de Tunisie, Rue Hédi Karray-2049 Ariana Tunis

**Keywords:** heritability, repeatability, breeding values, TDM, L305, dairy flocks, size, Tunisia

## Abstract

Genetic evaluation in dairy cattle has been commonly carried following the lactation model (L305) and the test-day model (TDM), the purpose of this study was to test the adjustment and the accuracy of these main models in relation to the size of the Tunisian dairy flocks while assessing the effect of genealogical data availability on both approaches. Data were obtained from the Tunisian official milk recording system and cows were classified in accordance to the flock sizes into eight groups. Genetic parameters and breeding values were estimated per size group for 305-days (L305) and daily milk yields (TDM) through two animal models and by using 3 pedigrees of different quality. Contemporary groups were defined as herd*calving year for L305 and as herd*control year for TDM. Genetic evaluation approaches were compared by connecting the different obtained results. Fixed factors were observed to be differently significant per group of flock size explaining a specific variance of the average milk yield and that small flocks are mostly affected by environmental factors. Using TDM and an equilibrated pedigree file, genetic parameters were higher, breeding values were fairly compared leading to a more objective ranking of the cows and a better illustration of genetic variabilities between the flock groups. Low genetic variability and significant contribution of unfavorable environmental conditions were observed within the Tunisian dairy flocks.

## INTRODUCTION

Animal selection has been emerged to objectively identify best genetic animal materials for procreating the subsequent generations to ensure long-term genetic improvement of the most important economic traits in livestock (Colleau and Régaldo, 2001; Weiner and Rouvier, 2009). The availability of genetic and statistic tools together with computer resources advancement have emerged different algorithmic and methodological approaches for livestock genetic evaluation which have facilitated identifying ideal genetic materials through an objective classification according to their most likely additive genetic values, known as breeding value (Colleau, 1996; Boichard and Brochard, 2012). Genetic evaluation procedure requires relevant phenotypic and pedigree records which define the efficiency of the analysis (Bonaiti et al., 1990). In dairy cattle, the lactation model (L305) was commonly used for genetic evaluation but since November 2002, most of dairy countries have adopted a more sophisticated methodology called the “Test Day model (TDM)” which directly analyzes daily milk records (Leclerc et *al*., 2009). In this context, the aims of this study have been defined in order to identify the genetic model which is better adjusted to the dairy flock performances in Tunisia by exploiting the two main analytical approaches in genetic evaluation of dairy cattle (L305 and TDM) through testing the pedigree quality effect.

## MATERIAL AND METHODS

### Data

Data were obtained from the Tunisian official milk recording system of the Agency of Livestock and Pasture (O.E.P) and included 1 531 953 test day (TD) records. Since the reliability of the genetic evaluation procedure is mainly based on the data quality all unacceptable records were excluded and flocks with less than 50 cows were not considered. After editing, 146 734 TD recorded between the 25^th^ and 360^th^ days in milk (DIM) were used and have corresponded to 20 523 lactations (L305). Data were obtained from 27 649 Holstein cows in their first, second and third parity, calved and controlled between 2006 and 2011 and belonging to 116 flocks which sizes have varied between 50 and 1712 according to which cows were classified into 8 groups. In the pedigree file, the mothers were identified for 11 726 cows, both parents were identified for 10 307 cows and 5 616 cows didn’t have any genealogical data. Two separated files were generated from the original one which included respectively only cows whose mothers were identified and cows whose both parents were identified. Data are described in Table I.

**Table I.** Characteristics of the retained records by herd size class

| Parameters | HS=50 | 50<HS<100 | 100≤HS<150 | 150≤HS<220 | 220≤HS<300 | 300≤HS<400 | 400≤HS≤600 | HS>600 | Total |
| --- | --- | --- | --- | --- | --- | --- | --- | --- | --- |
| Number of the total cows | 1 857 | 1 915 | 2 663 | 2 131 | 2 501 | 1 435 | 1 679 | 1 964 | 16 145 |
| Number of test-day records | 16 159 | 16 617 | 23 155 | 19 576 | 24 042 | 13 766 | 14 980 | 18 457 | 128 295 |
| Number of lactation records | 2 568 | 2 510 | 1 577 | 3 019 | 3 655 | 2 086 | 2 320 | 2 788 | 20 523 |
| Number of the mother know cows | 1 151 | 1 331 | 1 844 | 1 625 | 1 890 | 1 134 | 1 240 | 1 511 | 11 726 |
| Number of test-day records for the mother know cows | 10 104 | 11753 | 16 586 | 15 505 | 19 211 | 11 309 | 11 327 | 5 047 | 100 842 |
| Number of lactation records for the mother know cows | 1 606 | 1 762 | 2 553 | 2 361 | 2 889 | 1 708 | 1 750 | 2 249 | 16 878 |
| Number of the relative know cows | 901 | 1 145 | 1 552 | 1 427 | 1 811 | 1 075 | 1 034 | 1 362 | 10 307 |
| Number of test-day records for both relatives know cows | 7 756 | 10 127 | 13 832 | 13 632 | 18 316 | 10 745 | 9 507 | 13 356 | 97 271 |
| Number of lactation records for both relatives know cows | 1 231 | 1 526 | 2 128 | 2 081 | 2 759 | 1 623 | 1 466 | 1 995 | 14 809 |

HS= herd size

### Analysis

#### Estimation of the herd factor on the variation of the average milk yields

Evaluating the effect of the fixed factors on the milk yield average per day (TDMY) and per lactation (MY305) was performed per flock size group according to the following linear models (1 and 2):

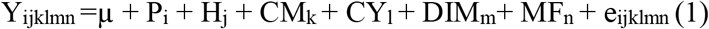

Where Y_ijklmn_ is the observed TDMY on the cow in parity i (i=1, 2, 3) that belongs to the herd of the j size (j = 2,…, 47) which was controlled in the month k (k=1, …, 12) during the year l (l = 2006,…, 2011) between the m days in milk (m = 1,…, 34) and milked with n frequency of milking (n = 1, 2, 3). µ = overall mean; P = the parity effect; CY = the control year effect; CM = the control month effect; CLM = the calving month effect; MF = the milking frequency effect; DIM = days in milk effect and e = the residual errors.

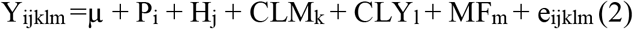

Where Y_ijklm_ is the observed MY305 on the cow in parity i (i = 1, 2, 3) that belongs to the j size herd (j = 2,…, 47) which was calved in the month k (k = 1,…, 12) during the year l (l = 2006,…, 2011) and milked with m frequency (m = 1, 2, 3). µ = overall mean; P = the parity effect; CLY = the calving year effect; CLM = the calving month effect; MF = the milking frequency effect and e = the residual errors.

The least squares solutions for daily and standard milk yield variation were obtained through the General Linear Model procedure (GLM) of the SAS program (Statistic Analysis System, 2003) and the contribution of each factor was calculated from the sums of square.

#### Genetic evaluation

The variance components of TDMY and MY305 within each group were obtained by two uni-trait animal models based on the method of the restricted maximum likehood. Contemporary groups were defined as herd*calving year for L305 and as herd*control year for TDM.

The Best Linear Unbiased Predector (BLUP) (Xu-Qing et al., 2008) applied to the general mixed linear animal model (Henderson, 1984; Gengler, 2000). The first animal model included the random additive genetic effect, the fixed effect of contemporary groups, the linear and quadratic regressions of the covariates herd, control year, control month, milking frequency, parity and DIM for TDM and for L305 the model included the random additive genetic effect, the fixed effect of contemporary groups, the linear and quadratic regressions of the covariates herd, calving year, calving month, milking frequency and parity. The model in matrix for both traits was as follow:

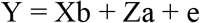

Where y is the vector which contained TDMY or MY305, b is vector of the fixed effect solutions, a is the vector of random genetic effect solutions, X and Z are matrix respectively relating the phenotype observations to the fixed and the genetic effects while e is the vector of random residual.

The second animal model included the random additive genetic effect, the permanent environment effect, the fixed effect of contemporary groups and the linear and quadratic regressions of the covariates herd, control year, control month, milking frequency, parity and DIM for TDM and for L305 it has included the random additive genetic effect, the permanent environment effect, the fixed effect of contemporary groups, the linear and quadratic regressions of the covariates herd, calving year, calving month, milking frequency and parity. The genetic and permanent environment effect model in matrix was as follow:

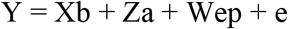

Where y is a vector which contained TDMY or MY305, b is the vector of the fixed effect solutions, a is the vector of the genetic random effect solution, ep is the vector of the permanent environment random effect solutions, X, Z and W are matrix relating phenotype observations to the different effects and e is the vector of residual.

The estimation of genetic parameters, the prediction of breeding values as well as the estimation of the permanent environment components was made per group and using each pedigree file by using the VCE 5.1 programme (Variance component estimator) (Kovach, 2002). Heritability (Verrier, 2001; Wiener et Rouvier, 2009) and repeatability (Minvielle, 1990; Bourdon, 1997) were estimated using the following formulas (3 and 4 respectively):

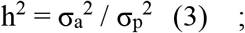

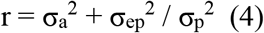

#### Comparison between TDM and L305

For the variance analysis, the estimation of the fixed factors contribution in the total observed variability in milk yield was compared between models through the coefficient of determination (R^2^), the F coefficient, the variation coefficient (CV) and the standard error coefficient. Regarding the genetic evaluation, we compared the genetic parameters and we calculated the rank coefficient of Spearman (ρ) between the predicted breeding values within each approach (L305 and TDM) for each animal model and each pedigree file within groups. Also, the genetic standard deviation distribution among groups were compared to illustrate the genetic variability obtained from the used animal model and pedigree file.

## RESULTS

### Estimation of the herd factor in the variance of TDMY and MY305

The herd factor compromises multiple fixed effects related to the husbandry and the environment conditions and which specifically characterize the breeding system of a given cow group. Thus, the herd factor represents a significant variation source and serves as an indicative element of the common non-genetic impacts that simultaneously influence average performances (Agabriel et al., 1990). Variance analysis have demonstrated that TDMY and MY305 have varied specifically within each flock group in accordance to the found coefficient of determination (table II) and to the different significance degrees of each common fixed factors (table III and IV) whose impact were found to be pronounced in the smallest herds which hold 50 cows. Besides we have noticed that the coefficients of determination revealed by the TDMY variance analyses were higher and the variance coefficients and F values were respectively lower than those found through the MY305 analyses.

**Table II.**
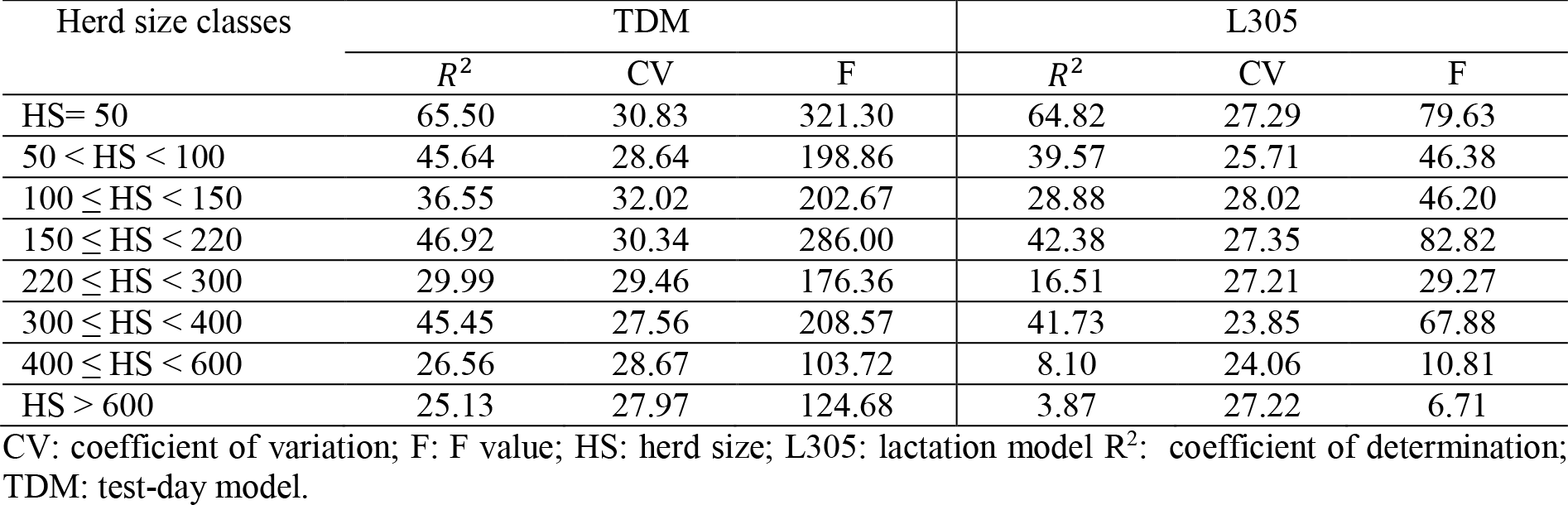
Determination coefficients of the variance analysis of MY305 and TDMY per herd size classes

| Herd size classes | TDM |  |  | L305 |  |  |
| --- | --- | --- | --- | --- | --- | --- |
| | $R^2$ | CV | F | $R^2$ | CV | F |
| HS= 50 | 65.50 | 30.83 | 321.30 | 64.82 | 27.29 | 79.63 |
| 50 < HS < 100 | 45.64 | 28.64 | 198.86 | 39.57 | 25.71 | 46.38 |
| 100 ≤ HS < 150 | 36.55 | 32.02 | 202.67 | 28.88 | 28.02 | 46.20 |
| 150 ≤ HS < 220 | 46.92 | 30.34 | 286.00 | 42.38 | 27.35 | 82.82 |
| 220 ≤ HS < 300 | 29.99 | 29.46 | 176.36 | 16.51 | 27.21 | 29.27 |
| 300 ≤ HS < 400 | 45.45 | 27.56 | 208.57 | 41.73 | 23.85 | 67.88 |
| 400 ≤ HS < 600 | 26.56 | 28.67 | 103.72 | 8.10 | 24.06 | 10.81 |
| HS > 600 | 25.13 | 27.97 | 124.68 | 3.87 | 27.22 | 6.71 |

CV: coefficient of variation; F: F value; HS: herd size; L305: lactation model $R^2$ : coefficient of determination; TDM: test-day model.

**Table III.** Source of variation of MY305 according to the herd size classes

| SV | N | FSC | ddl | SS | CM | F | Pr > F |
| --- | --- | --- | --- | --- | --- | --- | --- |
| Herd | 2919 | 1 | 46 | 1997207267 | 43417549 | 26.32 | *** |
|  | 2873 | 2 | 20 | 1588322583 | 79416129 | 32.09 | *** |
|  | 4131 | 3 | 16 | 1381561974 | 86347623 | 32.40 | *** |
|  | 3408 | 4 | 10 | 1939572986 | 193957299 | 87.36 | *** |
|  | 4172 | 5 | 9 | 1431066434 | 159007382 | 56.32 | *** |
|  | 2299 | 6 | 4 | 1280686105 | 320171526 | 138.48 | *** |
|  | 2694 | 7 | 2 | 433774960.8 | 216887480.4 | 64.17 | *** |
|  | 3180 | 8 | 1 | 1906974.8 | 1906974.8 | 0.39 | NS |
| Calving year | 2919 | 1 | 5 | 185210439 | 37042088 | 22.45 | *** |
|  | 2873 | 2 | 5 | 212138286 | 42427657 | 17.14 | *** |
|  | 4131 | 3 | 5 | 256243413 | 51248683 | 19.23 | *** |
|  | 3408 | 4 | 5 | 228082224 | 45616445 | 20.55 | *** |
|  | 4172 | 5 | 5 | 454528797 | 90905759 | 32.20 | *** |
|  | 2299 | 6 | 5 | 147958592 | 29591718 | 12.80 | *** |
|  | 2694 | 7 | 5 | 77653679.6 | 15530735.9 | 4.60 | NS |
|  | 3180 | 8 | 5 | 442837769.1 | 88567553.8 | 18.03 | *** |
| Calving month | 2919 | 1 | 11 | 37173296 | 3379391 | 2.05 | NS |
|  | 2873 | 2 | 11 | 142816248 | 12983295 | 5.25 | *** |
|  | 4131 | 3 | 11 | 64644815 | 5876801 | 2.21 | NS |
|  | 3408 | 4 | 11 | 72350050 | 6577277 | 2.96 | NS |
|  | 4172 | 5 | 11 | 171342960 | 15576633 | 5.52 | *** |
|  | 2299 | 6 | 11 | 49079077 | 4461734 | 1.93 | NS |
|  | 2694 | 7 | 11 | 86621044.3 | 7874640.4 | 2.33 | NS |
|  | 3180 | 8 | 11 | 89282115.9 | 8116556.0 | 1.65 | NS |
| Parity | 2919 | 1 | 2 | 88892893 | 44446446 | 26.94 | *** |
|  | 2873 | 2 | 2 | 96964489 | 48482244 | 19.59 | *** |
|  | 4131 | 3 | 2 | 6366645 | 3183323 | 1.19 | NS |
|  | 3408 | 4 | 2 | 196362477 | 98181239 | 44.22 | *** |
|  | 4172 | 5 | 2 | 13149790 | 6574895 | 2.33 | NS |
|  | 2299 | 6 | 2 | 42800512 | 21400256 | 9.26 | *** |
|  | 2694 | 7 | 2 | 85368438.7 | 42684219.3 | 12.63 | *** |
|  | 3180 | 8 | 2 | 48171921.1 | 24085960.5 | 4.90 | NS |
| Milking frequency | 2919 | 1 | 2 | 1107897976 | 553948988 | 335.76 | *** |
|  | 2873 | 2 | 2 | 654947278 | 327473639 | 132.31 | *** |
|  | 4131 | 3 | 2 | 345299497 | 172649748 | 64.79 | *** |
|  | 3408 | 4 | 2 | 276898009 | 138449005 | 62.36 | *** |
|  | 4172 | 5 | 1 | 27898591 | 27898591 | 9.88 | NS |
|  | 2299 | 6 | 2 | 705292760 | 352646380 | 152.52 | *** |
|  | 2694 | 7 | 2 | 131861592.8 | 65930796.4 | 19.51 | *** |
|  | 3180 | 8 | 1 | 26865591 | 37798591 | 7.86 | NS |

**SV** : source of variation; **N** : number of observations ; **FSC** : herd size class; **ddl** : degree of freedom ; **SS** : sum of the square ; **MS** : mean square; **F** : F value ; **Pr>F** : F probability; \*\*\* $P < 0,001$ ; **NS** : not significant at 5%.

**Table IV.** Source of variation of TDMY according to the herd size classes

| SV | N | FSC | ddl | SS | MS | F | Pr > F |
| --- | --- | --- | --- | --- | --- | --- | --- |
| Herd | 16679 | 1 | 46 | 104706,0766 | 2276,2191 | 103,26 | *** |
|  | 17121 | 2 | 20 | 80824,0513 | 4041,2026 | 126,92 | *** |
|  | 23987 | 3 | 16 | 65369,3454 | 4085,5841 | 112,11 | *** |
|  | 20121 | 4 | 10 | 109031,0979 | 10903,1098 | 381,42 | *** |
|  | 24761 | 5 | 9 | 91917,3277 | 10213,0364 | 296,28 | *** |
|  | 14072 | 6 | 4 | 72017,9689 | 18004,4922 | 573,78 | *** |
|  | 15538 | 7 | 2 | 47200,7373 | 23600,3686 | 477,06 | *** |
|  | 18996 | 8 | 1 | 366,1461 | 366,1461 | 6,81 | NS |
| Control year | 16679 | 1 | 5 | 2450,0988 | 490,0198 | 22,23 | *** |
|  | 17121 | 2 | 5 | 5194,3717 | 1038,8743 | 32,63 | *** |
|  | 23987 | 3 | 5 | 8720,7217 | 1744,1443 | 47,86 | *** |
|  | 20121 | 4 | 5 | 12428,1556 | 2485,6311 | 86,95 | *** |
|  | 24761 | 5 | 5 | 8578,7736 | 1715,7547 | 49,77 | *** |
|  | 14072 | 6 | 5 | 2086,1107 | 417,2221 | 13,30 | *** |
|  | 15538 | 7 | 5 | 1964,1364 | 392,8273 | 7,94 | *** |
|  | 18996 | 8 | 5 | 12685,0381 | 2537,0076 | 47,17 | *** |
| Control month | 16679 | 1 | 11 | 6578,0390 | 598,0035 | 27,13 | *** |
|  | 17121 | 2 | 11 | 7199,2592 | 654,4781 | 20,56 | *** |
|  | 23987 | 3 | 11 | 17543,3210 | 1594,8474 | 43,76 | *** |
|  | 20121 | 4 | 11 | 8981,1119 | 816,4647 | 28,56 | *** |
|  | 24761 | 5 | 11 | 9401,0992 | 854,6454 | 24,79 | *** |
|  | 14072 | 6 | 11 | 4877,5682 | 443,4153 | 14,13 | *** |
|  | 15538 | 7 | 11 | 6614,7440 | 601,3404 | 12,16 | *** |
|  | 18996 | 8 | 11 | 6736,1882 | 612,3807 | 11,39 | *** |
| parity | 16679 | 1 | 2 | 6629,2828 | 3314,6414 | 150,37 | *** |
|  | 17121 | 2 | 2 | 5187,7041 | 2593,8521 | 81,47 | *** |
|  | 23987 | 3 | 2 | 3523,5626 | 1761,7813 | 48,35 | *** |
|  | 20121 | 4 | 2 | 12230,8526 | 6115,4263 | 213,94 | *** |
|  | 24761 | 5 | 2 | 2991,7156 | 1495,8578 | 43,39 | *** |
|  | 14072 | 6 | 2 | 3183,4070 | 1591,7035 | 50,73 | *** |
|  | 15538 | 7 | 2 | 7838,0340 | 3919,0170 | 79,22 | *** |
|  | 18996 | 8 | 2 | 816,0017 | 408,0009 | 7,59 | NS |
| Milking frequency | 16679 | 1 | 2 | 128021,2673 | 64010,6336 | 2903,79 | *** |
|  | 17121 | 2 | 2 | 72607,2157 | 36303,6078 | 1140,20 | *** |
|  | 23987 | 3 | 2 | 40212,6419 | 20106,3210 | 551,74 | *** |
|  | 20121 | 4 | 2 | 26177,5173 | 13088,7587 | 457,88 | *** |
|  | 24761 | 5 | 1 | 1415,9110 | 1415,9110 | 41,08 | *** |
|  | 14072 | 6 | 2 | 60673,3369 | 30336,6685 | 966,79 | *** |
|  | 15538 | 7 | 2 | 5739,4708 | 2869,7354 | 58,01 | *** |
|  | 18996 | 8 | 1 | 1325,9320 | 1425,9320 | 38,06 | *** |
| Days in milk | 16679 | 1 | 32 | 81029,5650 | 2532,1739 | 114,87 | *** |
|  | 17121 | 2 | 32 | 124332,9268 | 3885,4040 | 122,03 | *** |
|  | 23987 | 3 | 32 | 162258,6065 | 5070,5815 | 139,14 | *** |
|  | 20121 | 4 | 32 | 166504,5173 | 5203,2662 | 182,03 | *** |
|  | 24761 | 5 | 32 | 249307,5480 | 7790,8609 | 226,01 | *** |
|  | 14072 | 6 | 32 | 121949,1470 | 3810,9108 | 121,45 | *** |
|  | 15538 | 7 | 32 | 205347,6236 | 6417,1132 | 129,72 | *** |
|  | 18996 | 8 | 32 | 311442,7204 | 9732,5850 | 180,96 | *** |

**SV** : source of variation; **N** : number of observations ; **FSC** : herd size class; **ddl** : degree of freedom ; **SS** : sum of the square ; **MS** : mean square; **F** : F value ; **Pr>F** : F probability; \*\*\* $P < 0,001$ ; NS : not significant at 5%.

### Genetic evaluation for milk yields

#### Random additive genetic animal model

Genetic merit has been considered as a specific additive effect for each cow and available performances relating to her and her relatives have been converged to estimate heritability for MY305 and for TDMY within flock groups (table V). Heritabilities have ranged respectively from 0.16 to 0.35 and from 0.37 to 0.51 for MY305 and TDMY similarly to the coefficients found in Tunisian Holstein cows (Rekik et al., 2006; Hammami et al., 2009) and to the widely found values which oscillate from 0.20 to 0.40 (Boujenane, 2005), 0.27 to 0.43 (Campus et al., 1994) and 0.18 à 0.51 (Van Tassell et al., 1997). These found coefficients have explained that milk yield is a moderate heritable trait as it has been reported by the several authors listed above. Besides, we have noticed that heritabilities which were estimated through TDM model were remarkably higher than those found under the L305 model. Additive genetic values have varied between −2351 kg and 2515 kg for MY305 and between −16 kg and 15 kg for TDMY within flock groups (table VI). These values have explained a low genetic variability within and between the groups which was further approved by the genetic standard deviation distributions between the groups (figure 1) and may be explained by the convergence of breeders in their choice of the genetic materials, as Agabriel et al. (1990) have reported. We have noticed that TDM has better demonstrated this trend (R^2^= 0.63) rather than L305 model (R^2^= 0.34).

**Table V.**
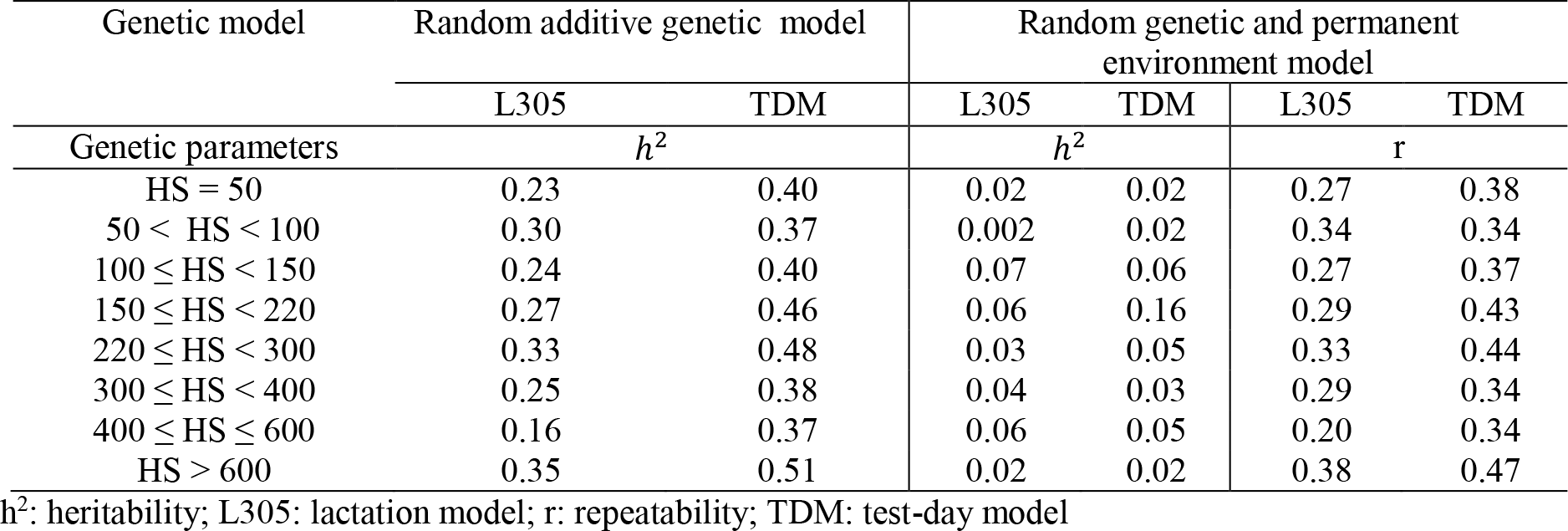
Genetic parameters for TDMY and MY305 according to herd size classes

**Table VI.** Genetic standard deviation under the random genetic animal model for TDMY and MY305 according to herd size classes

| Herd size classes | Number of cows | TDM |  |  | L305 |  |  |
| --- | --- | --- | --- | --- | --- | --- | --- |
|  |  | Standard deviation | Minimum | Maximum | Standard deviation | Minimum | Maximum |
| HS = 50 | 1 857 | 2.62 | -10.36 | 9.38 | 276.07 | -1222.66 | 1014.87 |
| 50 < HS < 100 | 1 915 | 2.92 | -13.12 | 10.33 | 405.20 | -1883.12 | 1404.51 |
| 100 ≤ HS < 150 | 2 663 | 3.33 | -12.32 | 14.55 | 375.07 | -1793.12 | 1509.36 |
| 150 ≤ HS < 220 | 2 131 | 3.31 | -12.69 | 14.07 | 390.60 | -1635.26 | 1373.09 |
| 220 ≤ HS < 300 | 2 501 | 3.65 | -15.29 | 15.81 | 518.95 | -2251.71 | 2515.33 |
| 300 ≤ HS < 400 | 1 435 | 2.95 | -10.57 | 10.16 | 376.00 | -1449.53 | 1386.42 |
| 400 ≤ HS ≤ 600 | 1 679 | 3.69 | -14.56 | 13.47 | 324.00 | -1475.12 | 1179.88 |
| HS > 600 | 1 964 | 4.68 | -16.63 | 13.93 | 684.64 | -2351.01 | 2549 |

**Figure 1.**
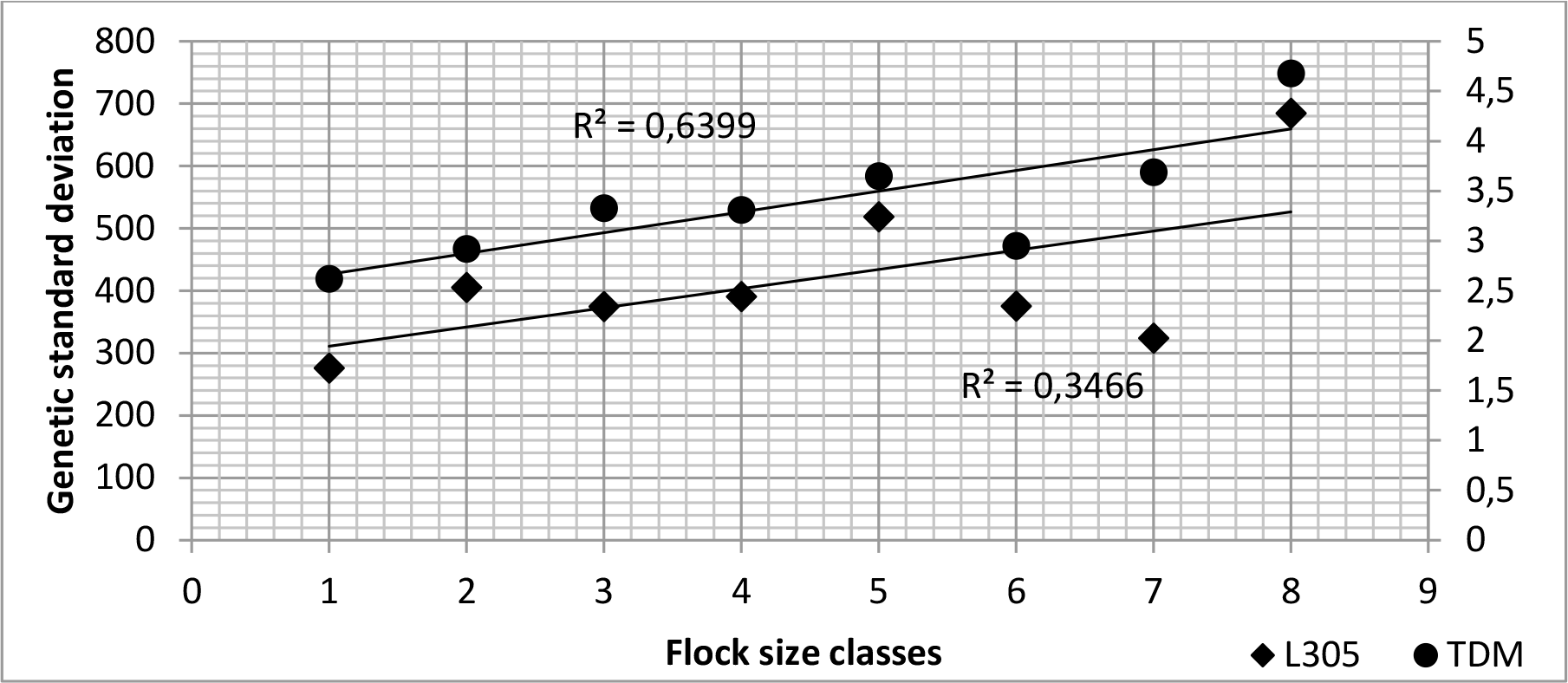
Distribution of the genetic standard deviation under TDM and L305 among flocks for the random genetic animal model

#### Random additive genetic animal model with permanent effect

Repetitive observations within a cow define the permanent environment effect which is a random specific non-transmissible value of the animal resulting from unidentified effects that are repeated each performance (Launay, 2015). Using this animal model, repeated performances were correlated thus heritability and repeatability were calculated (table V). Heritabilities were insignificant and have varied from 0.002 to 0.07 for MY305 and from 0.02 to 0.16 for TDMY. Repeatabilities have respectively varied between 0.20 and 0.38 and between 0.34 and 0.47. The found values were consistent with those reported in the Tunisian dairy cows by Rekik et al. (2006) and Hammami et al. (2009) with an average of about 0.39. We have found that repeatabilities and the variance component of the permanent environment (σ^2^_ep_) were higher than heretablities and additive genetic components (σ^2^_a_) which explains that the expression of genetic potential in Tunisian dairy cows seems to be influenced by an important contribution of additional non-identified environmental random factors as Laloë, 1992, and Hammami et al., 2009, have explained. Furthermore, we have found that the estimated heretablities with L305 are quite similar to those estimated with TDM for almost all the flock groups except for the group that counts from 150 to 220 cows. Repeatabilities were significantly different for all groups and were higher with TDM. Additive genetic values have varied between −607 kg and 518 kg for MY305 and between −4.62 kg and 5.89 kg for TDMY (table VII). Genetic standard deviations have respectively varied from 4.57 to 140.21 and from 0.18 to 1.28 while being remarkably lower than the permanent environment standard deviation coefficients which have respectively oscillated between 257.62 and 716.48 and between 2.12 and 4.42. In accordance to these values and referring to the distributions of the genetic standard deviations among group of flocks (figure 2), we have concluded that genetic diversity is low between the Tunisian dairy herds and that genotypes response to their specific environment is significantly divergent. This trend has been better illustrated with TDM (R^2^=0.06) rather than L305 (R^2^=0.28). Correlations between the predicted breeding values under TDM and L305 were positive explaining similar cow ranking according to their genetic merit; however, we have noticed that coefficient found within small herds, which size vary from 50 to 220 cows, were less significant (table VI).

**Table VII.** Genetic standard deviation under the random genetic and permanent environment animal model for TDMY and MY305 according to herd size classes

| Herd size classes | Number of cows | TDM |  |  | L305 |  |  |
| --- | --- | --- | --- | --- | --- | --- | --- |
|  |  | Standard deviation | Minimum | Maximum | Standard deviation | Minimum | Maximum |
| HS = 50 | 1 857 | 0.18 | -0.63 | 0.64 | 32.86 | -128.11 | 122.04 |
| 50 < HS < 100 | 1 915 | 0.20 | -1.28 | 4.57 | 4.57 | -16.11 | 16.20 |
| 100 ≤ HS < 150 | 2 663 | 0.65 | -2.83 | 2.79 | 130.51 | -607.53 | 518.33 |
| 150 ≤ HS < 220 | 2 131 | 1.28 | -4.62 | 5.89 | 106.97 | -373.76 | 336.94 |
| 220 ≤ HS < 300 | 2 501 | 0.59 | -1.70 | 2.00 | 88.13 | -241.30 | 287.51 |
| 300 ≤ HS < 400 | 1 435 | 0.37 | -1.36 | 1.25 | 93.05 | -298.28 | 349.64 |
| 400 ≤ HS ≤ 600 | 1 679 | 0.70 | -2.47 | 2.11 | 140.21 | -587.73 | 432.05 |
| HS > 600 | 1 964 | 0.37 | -1.13 | 0.94 | 75.66 | -238.06 | 230.12 |

**Figure 2.**
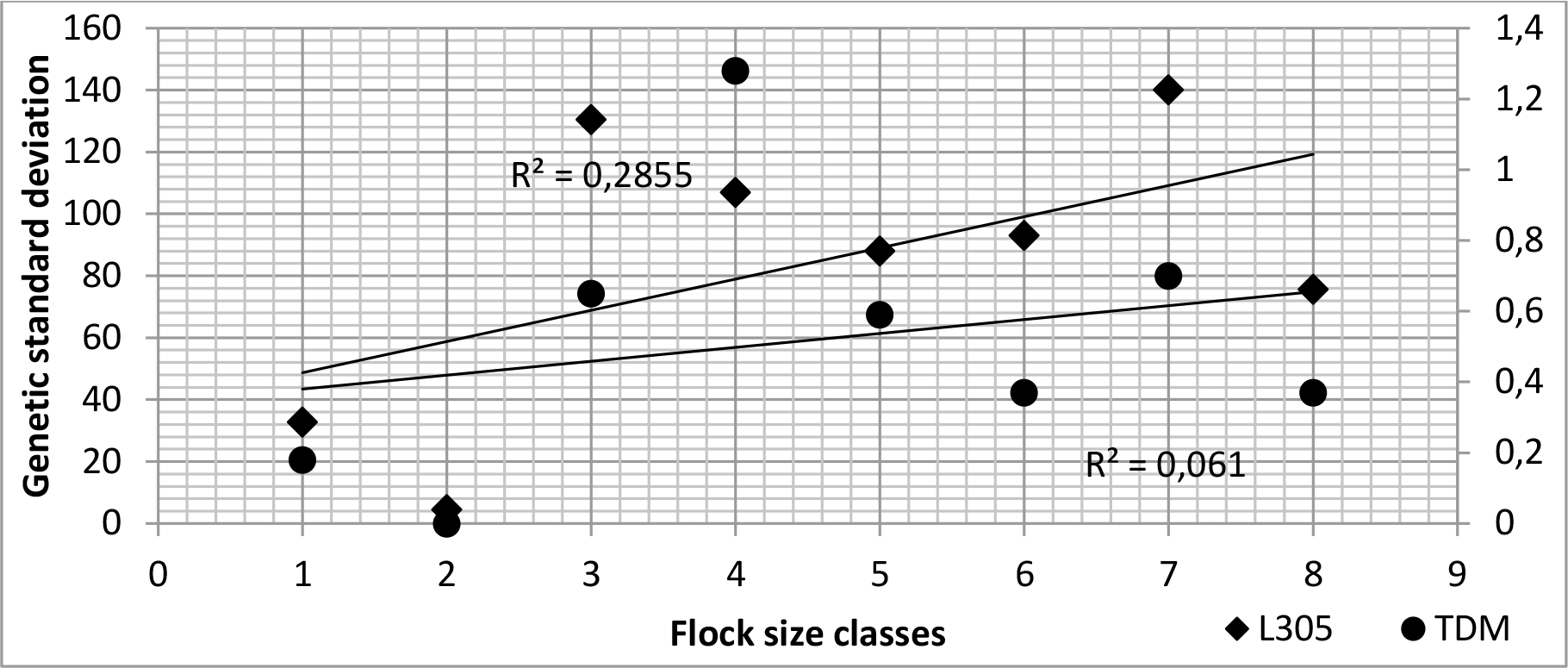
Distribution of the genetic standard deviation under TDM and L305 among flocks for the random genetic and permanent environment model

### Assessment of the pedigree quality

Generating the homogeneous pedigree files from the original one has led to reduce the number of observations as well as the number of animals for each flock category. Using these files, the genetic evaluation results within TDM and L305 were modified by systematically enhancing the genetic parameters estimates (table IX and X). Correlation coefficient between the predicted genetic values were positive and have fluctuated with L305 from 0.82 to 0.94 for the genetic additive animal model and from 0.10 and 0.99 for the animal model with both genetic and permanent environment effects. For TDM, coefficients have been higher and have respectively varied between 0.97 and 0.99 and between 0.52 and 0.99 (table VIII). The highest correlations were observed between the values which were predicted from equilibrate pedigrees. Distribution of genetic standard deviations in accordance to the used pedigree were also performed for both TDM and L305 (figure 3 and 4) and ameliorated trends were observed when the equilibrate pedigrees were used especially when considering the permanent environment effect; although the number of observations and animals have been reduced in both analyses.

**Table VIII.**
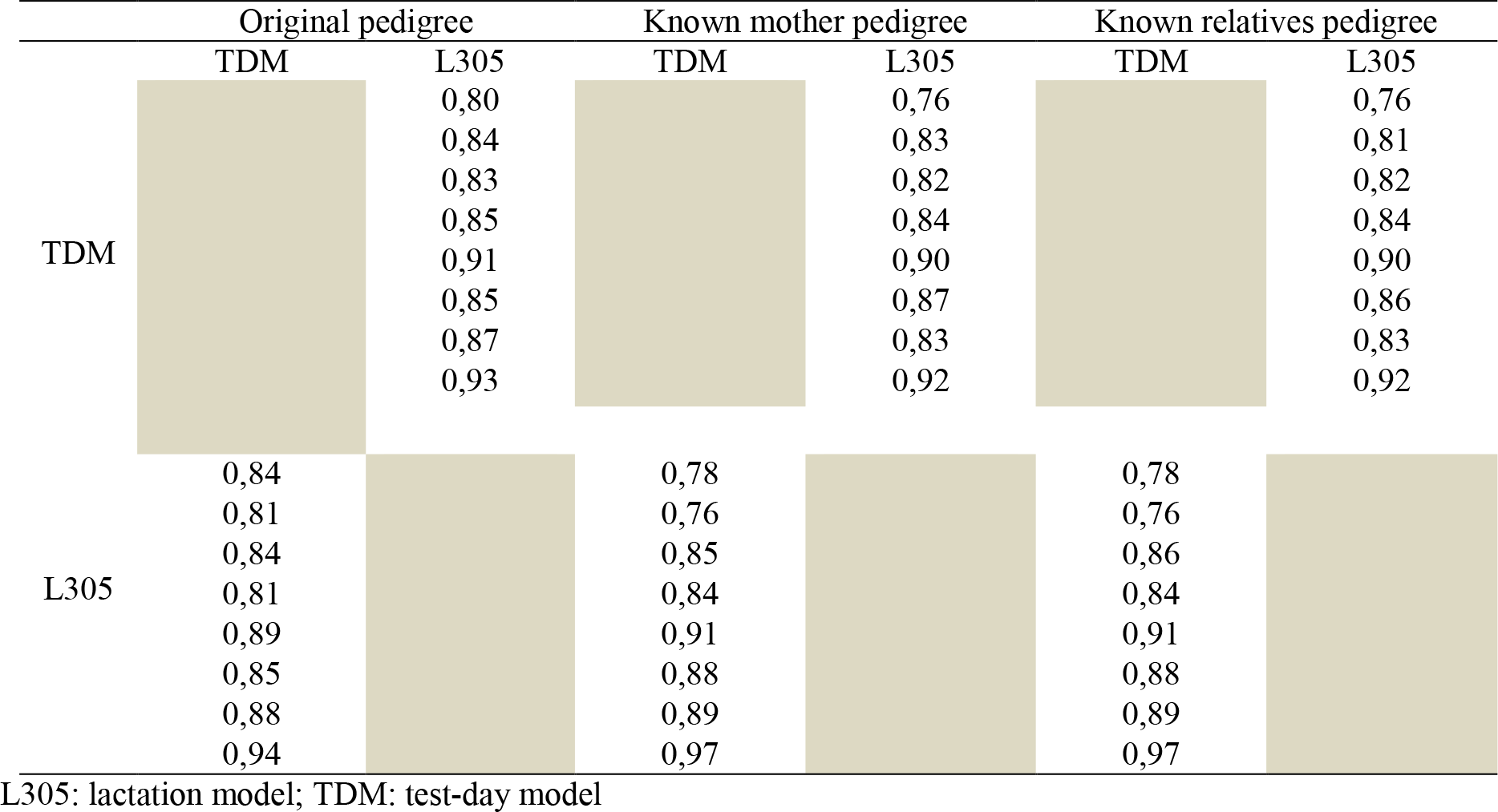
Ranking correlation coefficient according to the pedigree file under the random genetic animal model (above diagonals) and under the random genetic and permanent environment model (beyond diagonals).

**Table IX.** Heritability coefficient estimates under the random genetic animal model according to the pedigree file per herd size classes

| Genetic model |  | L305 |  |  | TDM |  |  |
| --- | --- | --- | --- | --- | --- | --- | --- |
| Pedigree files | P1 | P2 | P3 |  | P1 | P2 | P3 |
| HS = 50 | 0,23 | 0,21 | 0,21 |  | 0,40 | 0,36 | 0,37 |
| 50 < HS < 100 | 0,30 | 0,25 | 0,26 |  | 0,37 | 0,36 | 0,38 |
| 100 ≤ HS < 150 | 0,24 | 0,22 | 0,24 |  | 0,40 | 0,40 | 0,41 |
| 150 ≤ HS < 220 | 0,27 | 0,31 | 0,33 |  | 0,46 | 0,45 | 0,46 |
| 220 ≤ HS < 300 | 0,33 | 0,34 | 0,34 |  | 0,48 | 0,49 | 0,49 |
| 300 ≤ HS < 400 | 0,25 | 0,28 | 0,29 |  | 0,38 | 0,38 | 0,38 |
| 400 ≤ HS ≤ 600 | 0,16 | 0,15 | 0,18 |  | 0,37 | 0,37 | 0,38 |
| HS > 600 | 0,35 | 0,39 | 0,39 |  | 0,51 | 0,51 | 0,52 |

L305: lactation model; TDM: test-day model

**Table X.** Genetic parameters estimates under the random genetic and permanent environment model according to the pedigree file per herd size classes

| Genetic model |  | TDM |  |  |  |  |  | L305 |  |  |  |  |  |
| --- | --- | --- | --- | --- | --- | --- | --- | --- | --- | --- | --- | --- | --- |
| Genetic parameters | | $h^2$ | | | r | | | $h^2$ | | | r | | |
| Pedigree file | P1 | P2 | P3 | P1 | P2 | P3 |  | P1 | P2 | P3 | P1 | P2 | P3 |
| HS = 50 | 0.02 | 0.04 | 0.05 | 0.38 | 0.35 | 0.35 |  | 0.02 | 0.21 | 0.19 | 0.27 | 0.21 | 0.22 |
| 50 < HS < 100 | 0.02 | 0.04 | 0.05 | 0.34 | 0.34 | 0.35 |  | 0.002 | 0.01 | 0.01 | 0.34 | 0.34 | 0.34 |
| 100 ≤ HS < 150 | 0.06 | 0.10 | 0.09 | 0.37 | 0.37 | 0.37 |  | 0.07 | 0.05 | 0.04 | 0.27 | 0.25 | 0.26 |
| 150 ≤ HS < 220 | 0.16 | 0.22 | 0.30 | 0.43 | 0.32 | 0.44 |  | 0.06 | 0.22 | 0.24 | 0.29 | 0.32 | 0.33 |
| 220 ≤ HS < 300 | 0.05 | 0.02 | 0.03 | 0.44 | 0.33 | 0.43 |  | 0.03 | 0.01 | 0.02 | 0.33 | 0.33 | 0.33 |
| 300 ≤ HS < 400 | 0.03 | 0.04 | 0.02 | 0.34 | 0.30 | 0.34 |  | 0.04 | 0.05 | 0.06 | 0.29 | 0.30 | 0.31 |
| 400 ≤ HS ≤ 600 | 0.05 | 0.07 | 0.09 | 0.34 | 0.18 | 0.35 |  | 0.06 | 0.07 | 0.09 | 0.20 | 0.18 | 0.21 |
| HS > 600 | 0.02 | 0.08 | 0.09 | 0.47 | 0.46 | 0.46 |  | 0.02 | 0.11 | 0.11 | 0.38 | 0.39 | 0.39 |

$h^2$ : heritability; L305: lactation model; P1: original pedigree; P2: Known mother pedigree; P3: Known relatives pedigree; r: repeatability; TDM: test-day model

**Figure 3.**
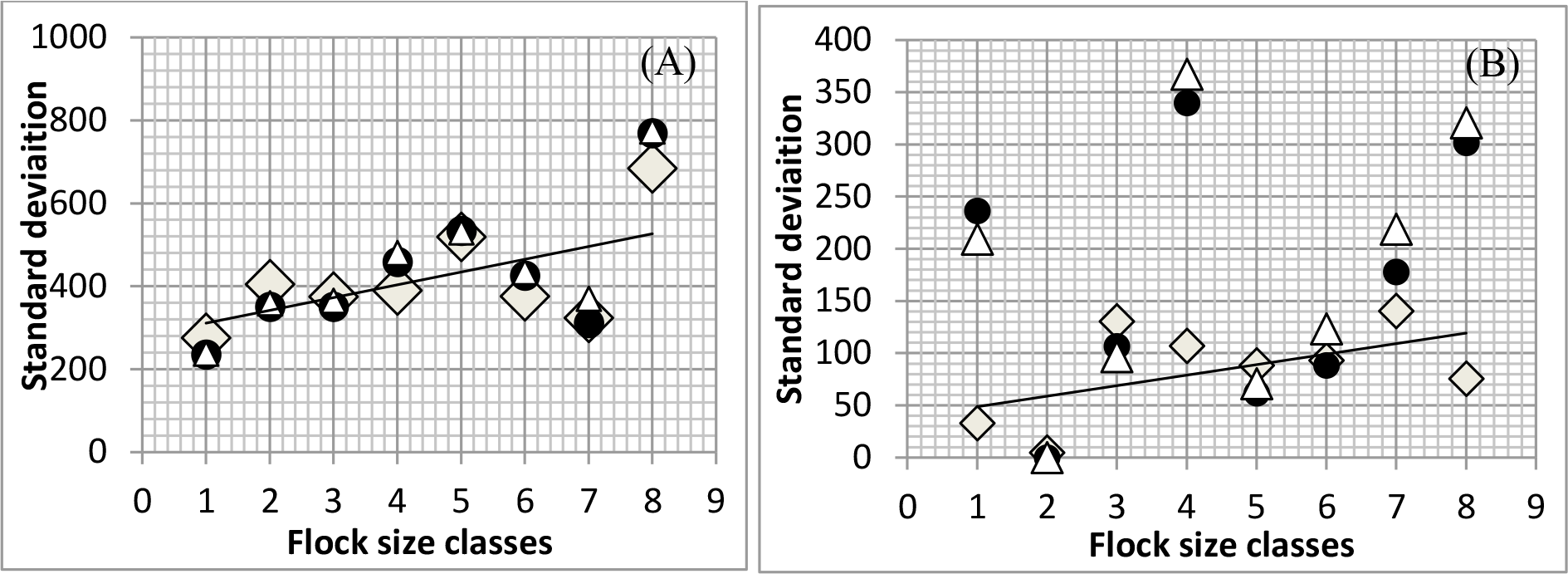
Distribution of the genetic standard deviation under L305 among flocks for the random genetic animal model (A) and for the random genetic and permanent environment model (B) according to the used pedigree

**Figure 4.**
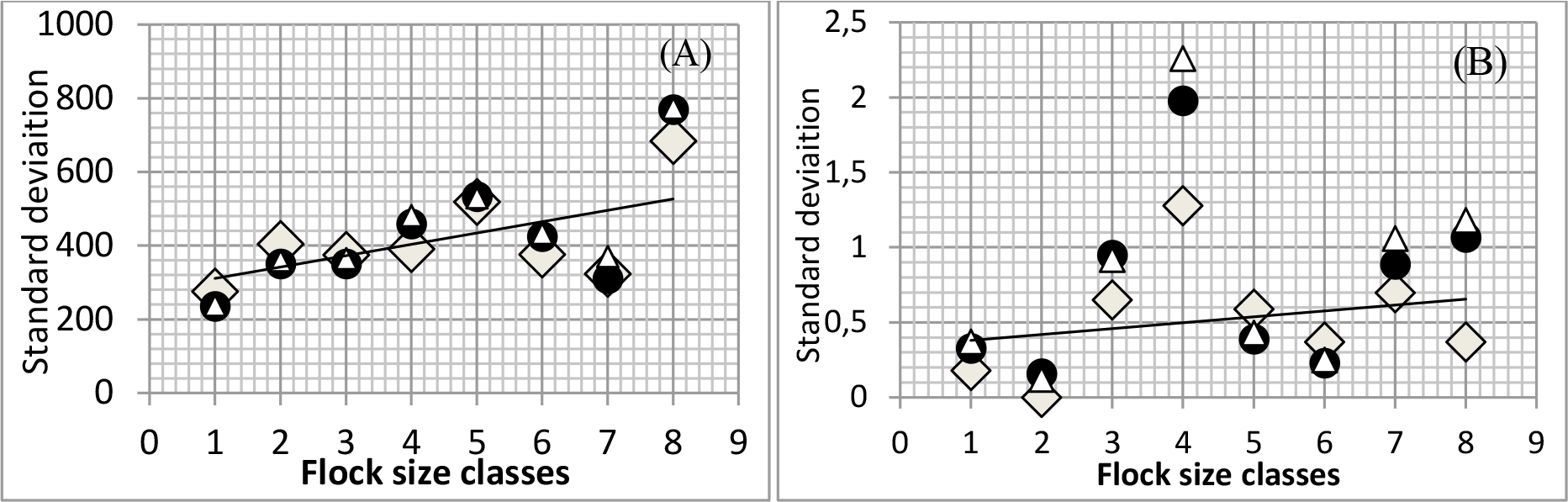
Distribution of the genetic standard deviation under TDM among flocks for the random genetic animal model (A) and for the random genetic and permanent environment model (B) according to the used pedigree files

## DISCUSSION

Variance analyses have demonstrated that fixed factors deeply contribute in variating the produced milk yield per day and per lactation. We also have found that each factor has his specific contribution within the different flock groups. We have noticed that dairy small herds in Tunisia are more affected by environmental factors comparing to medium and large herds and fixed effects seem to be much more involved in the expressed performances. In fact, it has been demonstrated that the herd factor in Tunisian dairy cows is a main factor that explains about 30% to 40% of the total observed variability in milk yield due to the differences that may exist between flocks regarding the geographic area of distribution, the relative climatic conditions, the breeding systems, the management facilities and all the relative related disadvantages (Accacha, 1985; Garrouri, 1986). In the same context Hamrouni et al. (2010) have found that the Tunisian dairy cows substantially interact with their environment as they found that different groups of genotypes respond in a particular way to different environmental circumstances of a given region and Rekik et al. (2003), have explained that remarkable differences in dairy sectors are as result of changes that might occur within the herd management and annual climatic conditions. Furthermore, our findings have proved oscillation in the yield level and its evolution between herds while the genetic materials are basically similar. This trend is in accordance with the study of Agabriel et al. (1990) which results have explained that observed heterogeneity in yields is due to a major contribution of the herd factor; as well as with the study of Jegou et al. (2005) who have reported that the herd factor reflects essentially the environmental conditions within a herd which define the Holstein cow productivity. Our results have explained that small dairy herds in Tunisia are raised under an inadequate management level and suffer from an important lack in facilities which, associated to dramatic climatic fluctuations, seem to greatly influence the genotype performances. This trend was more justified using the daily control analytical model which has better illustrated phenotype heterogeneities within and between herds. This observation underlines a better adjustment which is explained by minimum residual errors. Considering the permanent environment effect in both genetic evaluations, genetic variance components were reduced leading to a significant decrease in heretabilities and repeatabilities, which represent correlation between performances of the same animal. Furthermore, genetic standard deviations were found to be lower than permanent environment standard deviations reflecting moderate genetic diversity between herds and negative interaction between genotypes and their environment which have reduced breeding values of the different genotypes, as it has been demonstrated (Boujenane, 2005; Weiner and Rouvier, 2009). Our results have explained restricted genetic potential expression of dairy cows in Tunisia, as the animal model with direct genetic effect have shown higher heretabilities. Hence, the observed phenotype variabilities are due essentially to non-heritable factors and unidentified effects which are specific to each animal in every performance. Indeed, the contrasts between environments influence the adequate genetic merit expression of the desired phenotype which lead to a decrease in the corresponded heritability as it has been explained by Ollivier (1971) and Weiner and Rouvier (2009). Thus, the observed heterogeneity between herds is explained by the different responses of genotypes to environmental conditions which seem to be inadequate for milk yield especially in small herds. In the same context, Silvestre et al. (2006) have demonstrated that cows which are raised within a same region and under similar conditions but in different herds display different performances and that herd classification according to the management level could be very useful in identifying genetic variability; thus, it is then essential to correlate the genetic level of the animals with their environmental conditions (Hallais, 2013). The present study provides more affinity to the findings of Eddebbarh (1989) who has reported that, in the Mediterranean area, Holstein cows withstand harsh conditions such as thermal amplitudes, undernutrition and diseases against which they need to express extreme coping skills which enable them to express only the 1/3 of their genetic potential. Similar, Hammami et al. (2009) have highlighted that the major difficulties in genetic improvement of dairy cattle in many developing countries, including Tunisia, where small herds are predominate, are essentially related to environmental constraints and that the interaction between the genotype and the environment is a main key to develop sustainable breeding programs as long as the exporting countries of this breed need more knowledge about its environmental sensitivity within the importing countries to ensure genetic progress dissemination. This genetic trend found with our local data was more explained by the TDM suggesting that the L305 model is less accurate and precise due to significant random errors. The daily model TDM has shown more suitable to determine variability among herds as it considers specific environmental effects within each daily control rather than the whole lactation. Using the genetic animal model, cows were approximately ranked similarly according to their breeding values except for small herds. By including the permanent environment effect, highest correlations were observed within the large herds, which hold 600 cows and over, and were slightly lower for herds that contain about 150 to 400 cows. Lowest correlations were associated to small herds holding from 50 but not more than 100 cows. These results have explained that, in dairy cattle, TDM ensures better genetic evaluation analysis for milk records as it evaluates daily environmental factors. This essential advantage facilitates the use of a large data set and therefore helps to improve the accuracy of the genetic values estimation, especially for small herds, as it has been explained by Mayeres et al. (2004). To generate the balanced pedigree files, the number of cows has been respectively reduced from 16 145 to 11 726, while retaining only animals whose mothers are known, and to 10 307 by retaining only the known parent animals. Besides, the number of records was reduced from 128 295 to 100 842 to 97 271 test-day and from 20 523 to 16 878 and to 14 809 lactations. Using these files, the estimation quality has varied depending on the animal model within each evaluation procedure for the different classes.

Under the genetic animal model, heretablities were lower for L305 model in small herds have experienced decline in heretablities under the L305 model for both balanced pedigrees while this has been observed within TDM only when using the known mother pedigree whereas coefficients were observed to be higher for these classes with the known relatives pedigree except herds holding only 50 cows; regarding the remaining medium and large herds values have mostly been improved or held constant through the equilibrated files whether for L305 and for TDM.

Considering both genetic and permanent environment random effects, repeatabilities were observed to be reduced within small herds under L305 and TDM for both equilibrate pedigrees while heretablities were characterized by a complete different pattern according to which L305 have showed decreased values within small herds for both balanced pedigrees except within herds with 50 cows whose coefficients have been improved while under TDM all values have increased for all herds. Furthermore correlations were observed to be extremely significant just between the breeding values which were predicted from the balanced files and the distribution of genetic standard deviation between herds were proved to be more diversified also using these files especially by investigating the permanent environment effect and through the analysis of elementary daily controls. Oscillation of the results in accordance to availability of the genealogical information have proved that genetic evaluation procedure is essentially based on the relevance of these data by demonstrating that a bad quality pedigree which is marked by heterogeneity in the individuals identification and lack of information among the related animals may biased results especially when developing environmental effects as long as any individual without relatives known is assumed unrelated to the studied population. Thus balanced and accurate pedigrees are required to better detect the hereditary of the studied trait and to ensure fair comparison between the different individuals whose classification will be improved which will facilitate identifying the best genetic potentialities in terms of generations and jointly ameliorate exposure the genetic variability within populations, as Danchin-Burge and Moureaux, 2009, and Weiner and Rouvier, 2009, have reported.

## CONCLSION

In Tunisian Holstein herds, the herd factor has proved to be primarily responsible for the observed heterogeneity in the averages of daily and standard milk yield which was dictated to be a moderately transmissible trait. Low genetic components were revealed within the Tunisian dairy herds due to a lack of genetic variability either from deficiency in management and relatively harsh environmental conditions that define the adequate expression of the highest genetic potential of animals especially in small herds which seem to be subject to more pronounced effect. Inappropriate animal identification generates poor pedigree quality that disadvantages the adequate genetic variability determinism and the objective identification of the genetically interesting materials suggesting the improvement of milk recording operation in Tunisia. Analysis of daily controls ensures better quality of adjustment by illustrating more heterogeneity between performances and by providing less random errors; hence the milk yield is better described as well as the specific herd characteristics following a more detailed determination of the impact of a given environment. Therefore using TDM is recommended, specifically for small herds which will help to better simulate the impact of changes in breeding strategies and to enhance studying the genetic adaptability to the diverse livestock systems.

